# Selective Stabilization of HRAS2 i-Motif DNA by TMPyP4: A Multimodal Biophysical and Thermodynamic Investigation

**DOI:** 10.64898/2026.04.08.717182

**Authors:** Sagar Bag, Souvik Ghosal, Mangal Deep Burman, Erik Chorell, Sudipta Bhowmik

## Abstract

I-motif (iM) DNA structures, formed by cytosine-rich sequences, are increasingly acknowledged for their involvement in gene regulation, maintenance of genomic stability, and their emerging potential as therapeutic targets, particularly in cancer. Despite their biological relevance, the discovery of selective small-molecule probes that can specifically recognize and interact with iM DNA remains an ongoing challenge. In this study, we have used TMPyP4 and screened for its ability to bind various iM DNA constructs, including HRAS1, HRAS2, VEGF, CMYC, CKIT and H-Telo. Structure-activity relationship analyses revealed that specific substitution patterns conferred selectivity towards HRAS2 iM target. Comprehensive spectroscopic investigations, including UV-Vis absorption, steady-state and time-resolved fluorescence, and fluorescence anisotropy, uncovered key photophysical signatures of binding, including significant hypochromic and bathochromic shifts, enhanced fluorescence emission, and prolonged fluorescence lifetimes. Circular dichroism (CD),thermal denaturation (UV-melting) and thermodynamic investigations confirmed that TMPyP4 effectively stabilized the HRAS2 iM structures without disrupting their native topologies. Meanwhile, FT-IR spectroscopy revealed local structural rearrangements upon TMPyP4 binding, offering further evidence of molecular interaction. Collectively, these findings provide valuable insights into the molecular recognition of iM DNA by TMPyP4 and highlight its promise as both selective HRAS2 iM-binding agent and responsive fluorescent probe. This work lays a strong foundation for the development of novel tools for studying iM structures in biological systems and for designing future therapeutics targeting iM DNA in cancer and related diseases.

## INTRODUCTION

I-motifs (iMs) are dynamic, non-canonical DNA secondary structures that form within cytosine-rich sequences. Unlike the guanine-based Hoogsteen pairing found in G-quadruplexes, iM structures are stabilized by hemi-protonated cytosine–cytosine⁺ (C·C⁺) base pairs, which are uniquely dependent on acidic conditions for protonation to enable base pairing. This pH-sensitive requirement for cytosine protonation renders iMs more stable under slightly acidic environments [1]. The frequent presence of cytosine-rich tracts in promoter regions and telomeric DNA underscores the biological significance of iMs [2]. These structures have been implicated in crucial genomic processes, including transcriptional regulation, DNA replication and repair, centromere function, and the maintenance of telomere integrity [3]. In addition to their functional relevance in cellular biology, iMs demonstrate high responsiveness to environmental stimuli, lending themselves as promising elements in biosensing applications and nanobiotechnology [4]. Given their emerging role in modulating gene expression and genome dynamics, particularly in regions relevant to oncogenesis, iM structures are gaining attention as potential therapeutic targets in cancer and other disease contexts.

TMPyP4 (meso-tetra(N-methyl-4-pyridyl)porphyrin) is a cationic porphyrin compound [5] that has attracted significant attention in nucleic acid research because of its strong affinity for non-canonical DNA structures. Owing to its planar aromatic porphyrin core and positively charged pyridyl substituents [6], TMPyP4 can interact effectively with nucleic acids through π–π stacking and electrostatic interactions with the negatively charged DNA backbone. Although TMPyP4 has been extensively studied for its interaction with non-canonical DNA structures, most previous research has primarily focused on G-quadruplex DNA. Numerous studies have demonstrated that TMPyP4 can effectively bind and stabilize G-quadruplex structures present in the promoter regions of several oncogenes, thereby influencing gene expression and attracting attention for potential anticancer applications [7, 8]. In contrast, comparatively little attention has been given to the interaction of TMPyP4 with iM DNA structures. Although iMs are also found in the promoter regions of several cancer-associated genes and are believed to play important roles in gene regulation, systematic studies examining how TMPyP4 recognizes and stabilizes these cytosine-rich structures remain limited. Therefore, exploring the interaction between TMPyP4 and different iM DNA sequences is important for improving our understanding of ligand recognition of these structures and for expanding the potential applications of TMPyP4 in studying cancer-related iMs.

The main objective of this study was to investigate the interaction of TMPyP4 with several iM forming DNA sequences derived from cancer-associated genes. Another key goal was to determine whether TMPyP4 shows preferential binding and stabilization toward a particular iM structure. This study will contribute to this emerging field by exploring how TMPyP4, a well-known porphyrin compound, interacts with different iM DNA structures. By examining these interactions in detail, the work will provide insights into how TMPyP4 can selectively recognize particular iM sequences. Such knowledge may help in designing selective probes to detect iM structures in living systems and could also support the development of novel strategies to regulate oncogene activity, which is highly relevant in cancer research.

In this study, we examined the interaction of TMPyP4 with several iM forming DNA sequences originating from oncogene promoter regions and the human telomeric region. The cytosine-rich sequences investigated include HRAS1, HRAS2, CMYC, H-Telo, CKIT, and VEGF **(Table 1)**. To comprehensively evaluate these interactions, a series of spectroscopic and biophysical techniques were applied, including UV-Vis absorption spectroscopy, steady-state fluorescence measurements, fluorescence anisotropy, time-resolved fluorescence decay analysis, circular dichroism (CD) spectroscopy, thermal denaturation (UV-melting), analysis of thermodynamic parameters and Fourier transform infrared (FT-IR) spectroscopy. The results indicate that TMPyP4 shows a marked binding preference for the HRAS2 iM DNA structure compared with the other examined iM sequences. The interaction is supported by characteristic spectroscopic changes, such as pronounced hypochromic and bathochromic shifts in the absorption spectra, decreased fluorescence intensity, and increased fluorescence lifetimes, suggesting strong ligand–DNA association. Furthermore, CD and thermal melting studies reveal that TMPyP4 enhances the stability of the HRAS2 iM structure while preserving its native conformation. The thermodynamic analysis suggests that the interaction between TMPyP4 and HRAS2 iM DNA is primarily driven by enthalpy and occurs spontaneously. Complementary FT-IR analysis suggests subtle structural adjustments within the HRAS2 iM DNA framework upon TMPyP4 binding. Overall, these findings demonstrate the selective recognition and stabilization of HRAS2 iM DNA by TMPyP4, highlighting its potential application as a fluorescent probe and molecular tool for investigating iM structures in biologically relevant environments.

**Table 1:**
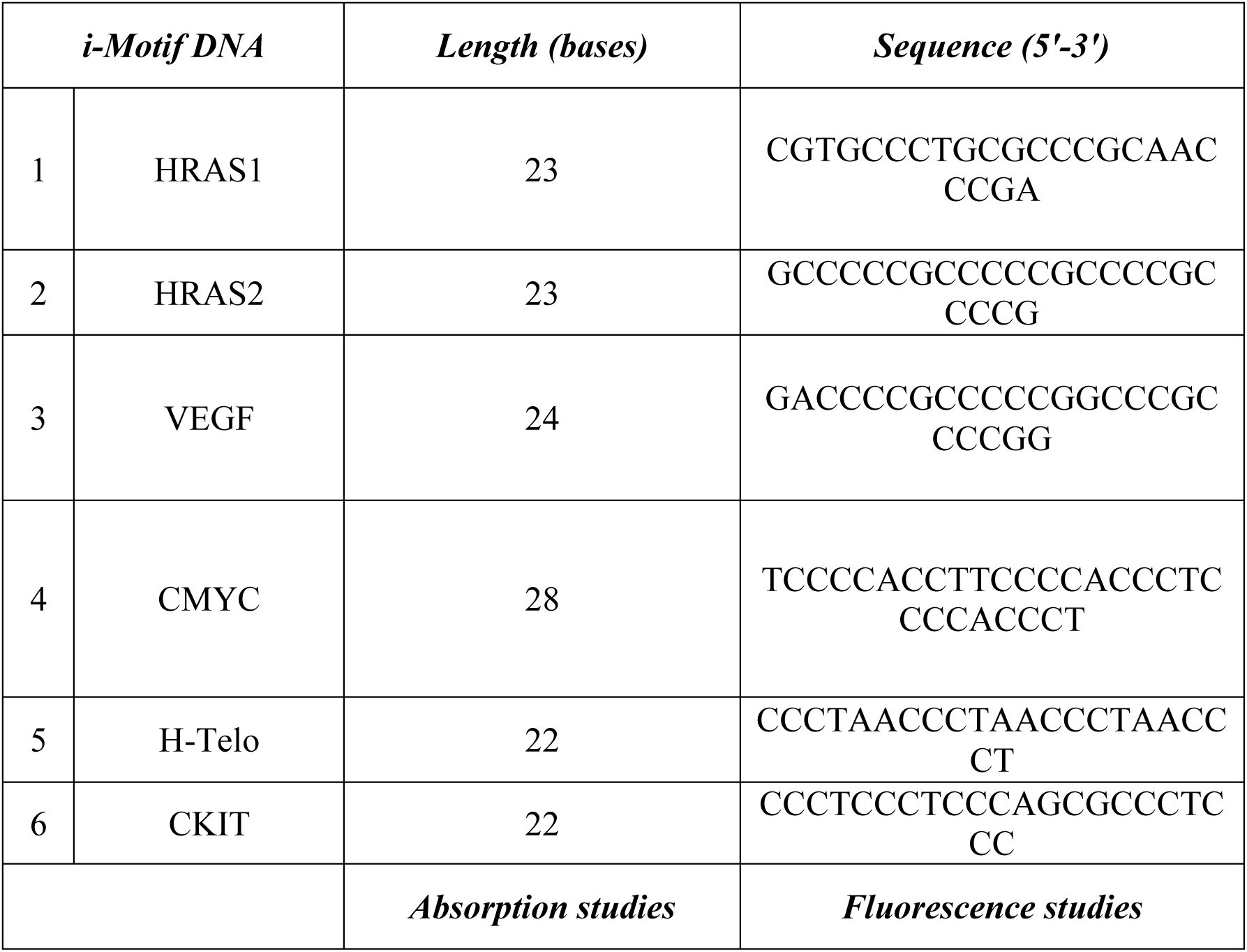
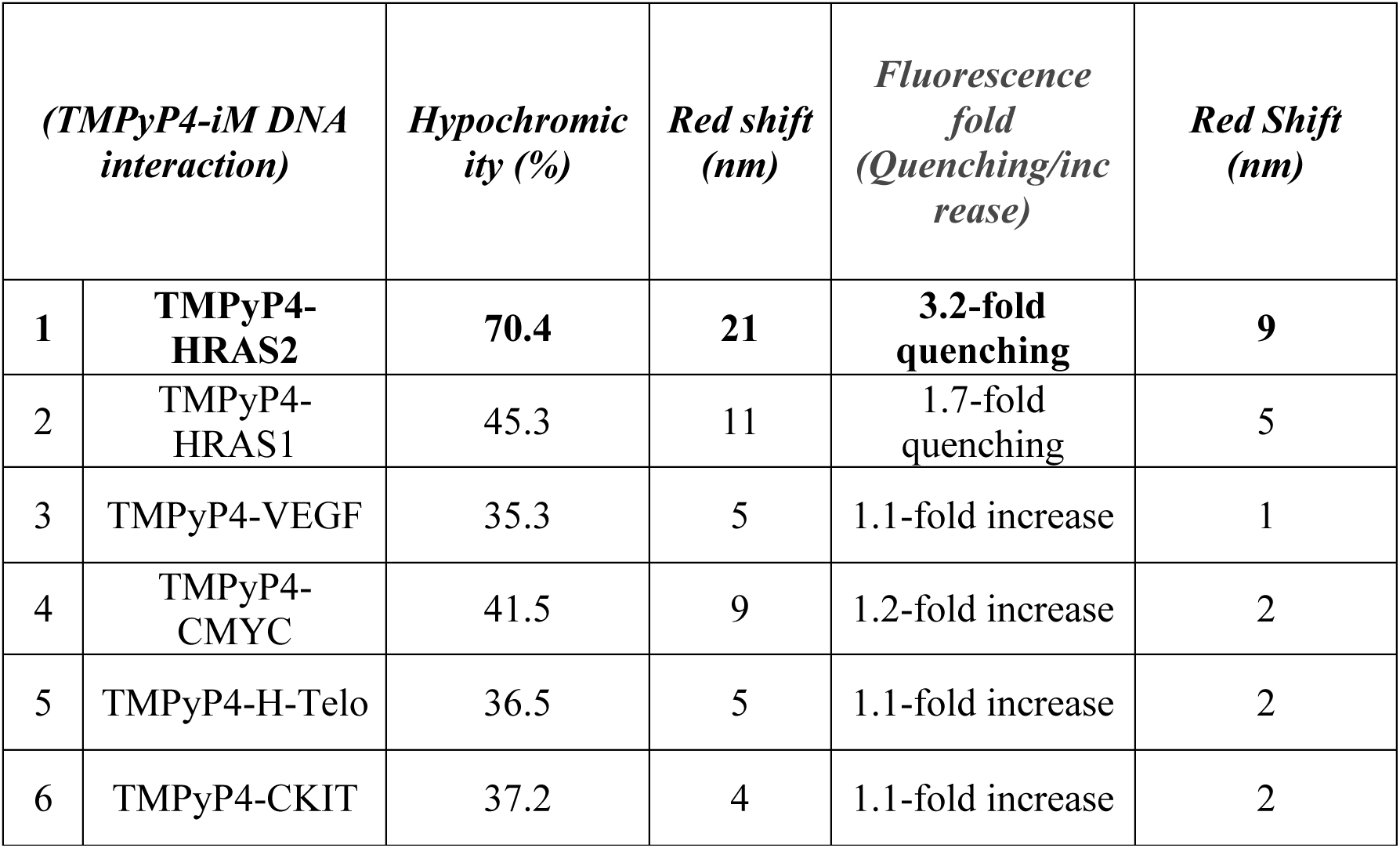
TMPyP4 and iM forming sequences used in this present investigation. Spectral parameters of interactions between TMPyP4 with different iM DNA (HRAS2, HRAS1, VEGF, CMYCH-Telo and CKIT) structures obtained from UV-Vis absorption and steady-state fluorescence studies.

## EXPERIMENTAL SECTION

### Materials, Oligonucleotides, and Stock Solution Preparation

TMPyP4 and several iM-forming DNA sequences, including HRAS1, HRAS2, VEGF, CMYC, CKIT and H-Telo were obtained from Sigma-Aldrich and used without any further chemical modifications. All solvents employed were of spectroscopic grade. To induce iM structure formation, a buffer solution containing 50 mM KCl, 10 mM KH₂PO₄, and 1 mM K₂EDTA was prepared at pH 5.4 and maintained at 25 °C. The oligonucleotides, after desalting and dissolution in double-distilled water, were stored at 4 °C. To assemble the iM DNA structures, the required quantity of each DNA was diluted into the respective buffers. The samples were then heated to 95 °C for 5 minutes to denature the strands, followed by gradual cooling to room temperature (25 °C), and subsequently kept at 4 °C overnight to ensure complete structural formation.

### UV-Vis Absorption Analysis

To investigate the preliminary binding interactions between TMPyP4 and various iM forming DNA structures, UV-Vis absorption spectroscopy was performed using a Hitachi UH5300 spectrophotometer. Measurements were carried out in quartz cuvettes with a 1 cm path length. During titration experiments, the concentration of TMPyP4 was maintained at 15 μM, while incremental amounts of iM DNA (ranging from 0 to 30 μM) were added to the solution. The experiments were conducted in buffer systems adjusted to pH 5.4 at 25 °C. Spectral data were recorded at a scanning rate of 400 nm/min, with measurements taken at 1 nm intervals.

### Steady-State Fluorescence Spectroscopy and Binding Constant Calculation

Fluorescence emission studies of TMPyP4 and iM DNAs were carried out using a Biobase BK-F96Pro fluorescence spectrometer equipped with a 1 cm path-length quartz cuvette. In these titration experiments, the concentration of TMPyP4 was fixed at 10 μM, while increasing concentrations (0–30 μM) of various iM-forming DNA sequences were introduced. Measurements were conducted in buffer systems maintained at pH 5.4 at 25 °C. The excitation and emission slit widths were set to 5 nm and 10 nm, respectively. Emission spectra were recorded at an excitation wavelength of 365 nm, while emission wavelengths were scanned over the relevant spectral range at a rate of 200 nm/min, with a response time of 0.5 seconds.

Binding constants (K_b_) were determined using a modified Benesi–Hildebrand approach based on the fluorescence intensity data [9]. This method relies on fluorescence spectroscopy, as it is both sensitive and accurate and works well even with low sample volumes. The K_b_ values were obtained using the Benesi–Hildebrand equation by relating fluorescence changes to iM DNA concentration. The relationship was modelled using the equation:

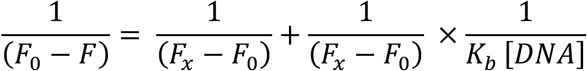

Here, F_0_= fluorescence intensity in the absence of iM DNA; F= fluorescence intensity at intermediate DNA concentrations; and F_X_= fluorescence intensity at saturation respectively. A linear plot of 1/F_0_-F against 1/[iM DNA] was used to extract the binding constant (K_b_) from the ratio of intercept to slope.

### Steady-State Fluorescence Anisotropy Measurements

Steady-state fluorescence anisotropy measurements were conducted using a Horiba Fluorolog spectrometer equipped with polarization filters. Samples were placed in 1 cm path-length Suprasil quartz cuvettes. Emission spectra for TMPyP4 were recorded over the 370–650 nm wavelength range, using an excitation wavelength of 365 nm. Both excitation and emission slits were set to 5 nm. All measurements were carried out in an experimental buffer system maintained at pH 5.4 and 25 °C.

Anisotropy values (r) were calculated using the following equation:

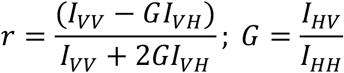

Here, I_VV_, I_VH_, I_HV_, and I_HH_ correspond to fluorescence intensities with the polarizers oriented at (excitation, emission): (0°, 0°), (0°, 90°), (90°, 0°), and (90°, 90°), respectively. The G-factor compensates for detector sensitivity discrepancies in perpendicular and parallel detection channels. Fluorescence anisotropy experiments were performed for 10 μM solutions of TMPyP4, both in absence and in the presence of 10 μM and 20 μM concentrations of HRAS2 and HRAS1 iM-forming DNA sequences. Each anisotropy value reported represents the mean of ten independent measurements.

### Time-Resolved Fluorescence Decay Measurements

Fluorescence lifetime studies were carried out using a Horiba Scientific DD-375L Fluorolog spectrometer equipped with time-correlated single-photon counting (TCSPC) capabilities. A 365 nm laser diode served as the excitation source for probing the TMPyP4. The instrument response function (IRF) had a full width at half maximum (FWHM) of 108 ps. To enhance the signal-to-noise ratio, 5000 photon counts were accumulated at the peak of the decay curve. Emission measurements were recorded using 5 nm slit widths, and an emission monochromator was used to suppress scattered light and isolate fluorescence signals.

Fluorescence decays were monitored at 520 nm for TMPyP4. The decay profiles were analysed using a multi-exponential fitting model in the DAS-6 (Decay Analysis Software) package:

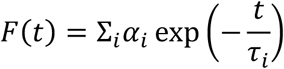

In this equation, F(t) is the fluorescence intensity at time t, while αi and τi represent the amplitude (pre-exponential factor) and lifetime of the i_th_ component. Fit quality was assessed using the chi-square (χ2) metric, as well as visual inspection of the residuals. The mean fluorescence lifetime (𝜏_*avg*_) was calculated using the following weighted equation:

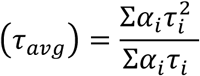

Fluorescence lifetime measurements were performed for 10 μM solutions of TMPyP4, both in the absence and presence of 10 μM and 20 μM of iM-forming DNA sequences HRAS2, and HRAS1. All experiments were conducted in the experimental buffer at pH 5.4 and 25 °C.

### Circular Dichroism (CD) Spectroscopy

CD spectra were recorded at 25 °C using a J-1500 spectropolarimeter (Jasco International Co. Ltd) with a 1 mm path-length quartz cuvette. Each spectrum represented the average of three independent scans, collected over the wavelength range of 200–350 nm at a scanning rate of 100 nm/min. The concentration of iM-forming DNA sequences—HRAS2 and HRAS1—was maintained at 20 μM, while the concentrations of TMPyP4 were varied (20, 40, 60, 80, and 100 μM). All measurements were performed in a buffer system adjusted to pH 5.4 and maintained at 25 °C. Buffer-only spectra were subtracted to account for background signals. Data analysis and curve fitting were performed using OriginPro 8 software.

### Thermal Denaturation (UV Melting) Assay and Thermodynamic Parameters Calculation

Thermal melting experiments were conducted using a Hitachi UH5300 spectrophotometer, coupled with a Julabo F12 temperature control system and a 3J1-0104 water-jacketed cuvette holder. The melting temperature (T_m_) was obtained from the UV melting profile by identifying the temperature at which the structural transition reaches its midpoint. In parallel, derivative analysis of the absorbance curve was used to confirm the T_m_ values. A quartz cuvette with a 1 cm path length was used for all measurements. iM DNA samples (10 μM HRAS2 and HRAS1) were monitored at 295 nm while being heated from 30 °C to 98 °C at a rate of 0.5 °C/min. Experiments were performed both in the absence and presence of 20 μM of TMPyP4. The thermal transition data were normalized and analysed using OriginPro 8 to determine melting behaviours. All measurements were carried out in the experimental buffer system (pH 5.4, 25 °C).

The thermodynamic parameters were estimated from the UV melting experiments. The melting curves were first normalized and then analysed through curve fitting using Kaleida Graph (Synergy Software) along with Origin 2019b. From this analysis, values of ΔH, ΔS, and ΔG were obtained for the formation of the TMPyP4–HRAS2 iM DNA complex. The Gibbs free energy (ΔG) was then determined using the standard thermodynamic relationship, where ΔG is given by the difference between enthalpy and the temperature-scaled entropy term.

### Fourier Transform Infrared Spectroscopy (FT-IR)

The interaction of TMPyP4 with HRAS2 and HRAS1 iM DNA structures was investigated using Fourier transform infrared (FT-IR) spectroscopy. Distinct vibrational peaks in the IR spectra serve as molecular fingerprints, enabling differentiation among various iM conformations. FT-IR spectra of HRAS2, and HRAS1 iM DNA—both in the absence and presence of TMPyP4—were recorded using a Spotlight 400 FT-IR spectrometer (PerkinElmer, UK) at room temperature (25 °C).

Each spectrum was acquired by averaging over 120 scans at a spectral resolution of 2 cm⁻¹. A blank attenuated total reflectance (ATR) plate spectrum was used as the reference background. Baseline correction and spectral analysis within the 800–2500 cm⁻¹ region were performed using the Opus 7.5 software package (Bruker Optics). All measurements were conducted on liquid samples under identical conditions to ensure consistency.

### Statistical Investigation

All experimental data were analysed using OriginPro 8 software. Mean values and standard deviations (± SD) were calculated for each dataset. Statistical significance was assessed using one-way ANOVA followed by Tukey’s post-hoc test when appropriate. A p-value of less than 0.05 (p < 0.05) was considered statistically significant. Binding constants (K_b_) and thermodynamic parameters were determined through nonlinear regression analysis, with goodness-of-fit evaluated using R² values. All experiments were performed in triplicate to ensure reproducibility and consistency. Results are reported as mean ± SD, with statistical significance indicated in figure legends and accompanying tables for clear data interpretation. Error bars representing standard deviations are included in all relevant graphical presentations.

## RESULTS AND DISCUSSIONS

### UV-Vis Absorption Analysis of TMPyP4 with iM DNA Structures

To obtain a preliminary understanding of the binding behaviour, UV–Vis absorption spectra of TMPyP4 were monitored with and without the addition of HRAS1, HRAS2, VEGF, CMYC, H-Telo, CKIT, BCL2 iM DNA constructs. To gain insight into the influence of iM DNA binding, UV–Vis absorption spectra were recorded. The observed reduction in absorbance together with a red shift in the absorption maximum suggests a change in the local environment of TMPyP4, likely associated with differences in polarity upon interaction.

Following extensive screening across different iM DNA sequences, TMPyP4 was observed to interact more strongly with the HRAS2 iM compared to other iMs **(Figure 1 and Figure S1)**. In the case of the TMPyP4–HRAS2 iM complex, a significant hypochromic effect of 70.4 % was observed, accompanied by a 21 nm red shift (bathochromic shift) in the absorption maximum. These spectral alterations suggest a change in the polarity of the local environment surrounding the TMPyP4 molecule upon its interaction with the HRAS2 iM DNA structure **(Figure 1**, **Table 1).** Collectively, this observation highlights the preferential binding of TMPyP4 to hydrophobic regions within the folded architectures of HRAS2 iM DNA. The binding appears to shield these compounds from the surrounding polar aqueous medium, supporting their utility as selective fluorescent probes for iM DNA detection.

**Figure 1.**
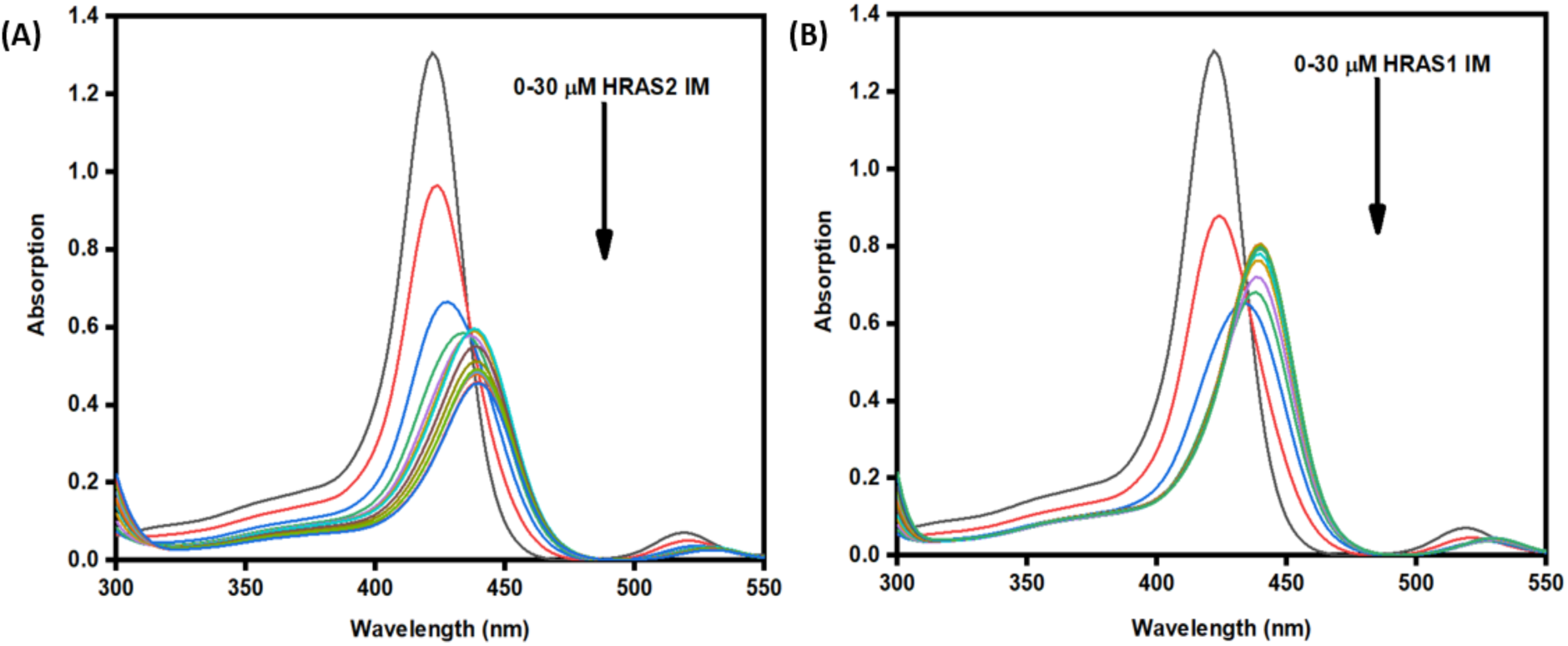
UV-Absorption spectra of TMPyP4 (15 μM) in the absence and presence of successive additions of HRAS2 (A) and HRAS1 (B) IM DNA (0-30 μM).

Typically, strong π-stacking interactions between molecules and nucleic acid bases cause major spectral changes, whereas weaker external interactions—such as binding in grooves or loop regions—are associated with minimal red shifts (Δλ < 8 nm) [10,11, 12]. The pronounced bathochromic and hypochromic effects observed in this study indicate that TMPyP4 interacts with HRAS2 iM DNA through a strong stacking-based binding mode rather than surface or groove association.

### Steady-state Spectrofluorometric Analyses

The addition of HRAS2 iM DNA caused a marked quenching of fluorescence emission from TMPyP4, confirming the formation of stable complexes. Specifically, TMPyP4 exhibited high selectivity for the HRAS2 iM DNA structure, with a 3.2-fold reduction in fluorescence intensity and a 9 nm red shift in emission **(Figure 2A**, **Table 1)**. These fluorescence shifts further support the distinct and selective interactions between the compounds and their respective iM DNA targets. No appreciable change in fluorescence intensity or emission maximum was observed upon the addition of HRAS1, CMYC, VEGF, CKIT and H-Telo iMs to TMPyP4 **(Figure S2)**. These findings are corroborated by UV-Vis absorption spectroscopy.

**Figure 2.**
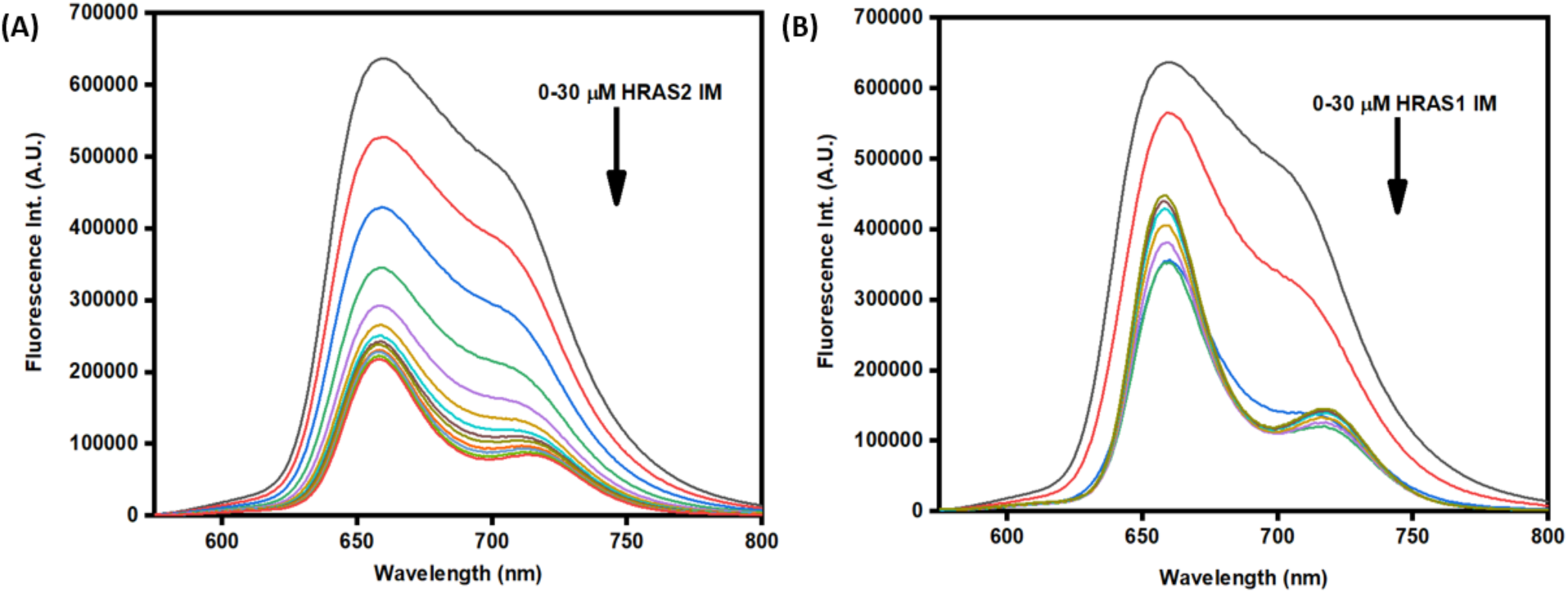
Upper panel: Fluorescence emission spectra of TMPyP4 (10 μM) in the absence and presence of successive additions of HRAS2 (A) and HRAS1 (B) IM DNA (30 μM).

A progressive decrease in the fluorescence intensity of TMPyP4 was observed as the concentration of HRAS2 iM DNA increased, suggesting fluorescence quenching due to complex formation. This quenching likely results from alterations in the local electronic environment of the TMPyP4 molecules upon interaction with the HRAS2 iM DNA. Binding to the HRAS2 iM DNA may restrict the molecules’ rotational and vibrational motions, thereby favour non-radiative relaxation pathways and reduce fluorescence emission. Additionally, the insertion of TMPyP4into hydrophobic or sterically constrained regions of the HRAS2 iM DNA may contribute π–π stacking interactions between the planar aromatic rings of the TMPyP4 and the nucleobases of HRAS2 iM DNA can further promote energy or electron transfer, intensifying the quenching effect. The observed fluorescence quenching in this case is better explained by changes in excited-state dynamics instead of just reduced molecular motion. While a more rigid environment can enhance emission in some systems, the results here suggest that non-radiative pathways, possibly involving interactions with the HRAS2 iM DNA bases, play an important role. Collectively, these fluorescence changes support the formation of stable, specific TMPyP4–HRAS2 iM DNA complex.

### Binding Affinity Determination Through Fluorescence Titration

To evaluate the binding affinity of TMPyP4 toward HRAS2 iM DNA structures, fluorescence titration experiments were conducted. Changes in fluorescence intensity as a function of increasing HRAS2 iM DNA concentrations were analysed using the modified Benesi–Hildebrand method [12]. The calculated binding constants (K_b_) revealed that TMPyP4 has a markedly higher affinity for the HRAS2 iM structure compared to HRAS1 iM DNA. The K_b_ value for the TMPyP4–HRAS2 iM complex was determined to be 5.67×10^5^ **(Figure 3A)**, while the TMPyP4–HRAS1 iM DNA interaction yielded a lower binding constant of 10.2×10^4^ **(Figure 3B)**.

**Figure 3.**
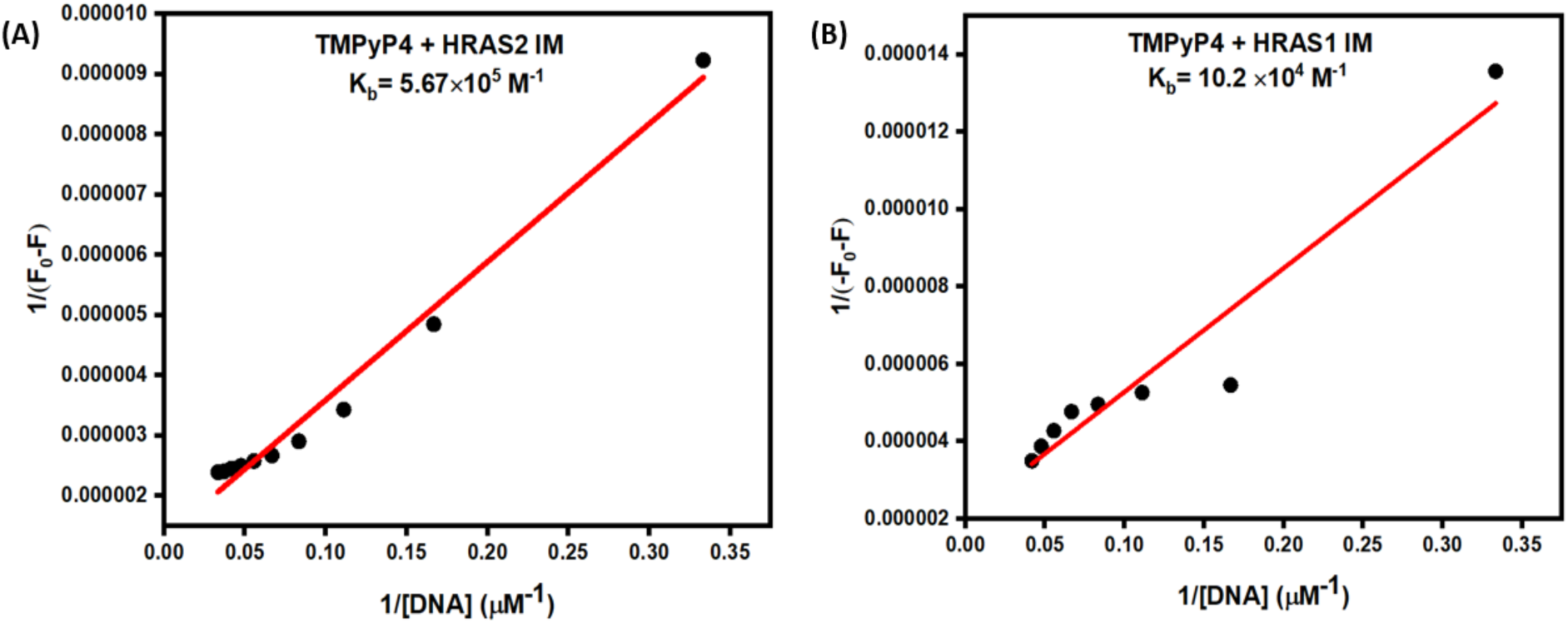
Modified Benesi-Hildebrand double reciprocal plots for determination of binding constant between TMPyP4 with HRAS2 IM (A) and HRAS1 (B) IM DNA.

Although both UV-Vis absorption and fluorescence spectroscopy are commonly employed to investigate ligand–DNA interactions, fluorescence spectroscopy was chosen in this study for determining binding constants due to its enhanced sensitivity and specificity. TMPyP4 possess strong intrinsic fluorescence, making them particularly suitable for fluorescence-based assays. This technique offers a significant advantage in detecting subtle emission changes induced by binding events, which might be difficult to discern through absorbance measurements, especially at low concentrations of either DNA or ligand.

Furthermore, fluorescence spectroscopy generally yields a higher signal-to-noise ratio and greater accuracy, particularly in systems where overlapping absorbance bands could complicate interpretation. Therefore, fluorescence titration was selected as the preferred method for characterizing the interactions between TMPyP4 and the HRAS2 iM DNA structures. Consistent with this approach, these fluorescence results agree well with the UV-Vis data and further support the selective binding of TMPyP4 to HRAS2 iM DNA. Based on the magnitude of the binding constants and the observed spectral responses, such as distinct bathochromic/blue shifts and fluorescence quenching/light-up effects, TMPyP4 was further investigated as potential molecular probe targeting HRAS2 iM DNA structure.

### Steady-State Fluorescence Anisotropy Analysis

Steady-state fluorescence anisotropy offers a sensitive approach for assessing the rotational dynamics of small molecules within macromolecular environments, providing direct insight into the degree of motional restriction imposed by binding interactions [13,14]. To further probe and substantiate the interactions of TMPyP4 with HRAS2 iM DNA structures, anisotropy measurements were performed **(Figure 4A–4B**; **Table 2**).

**Figure 4.**
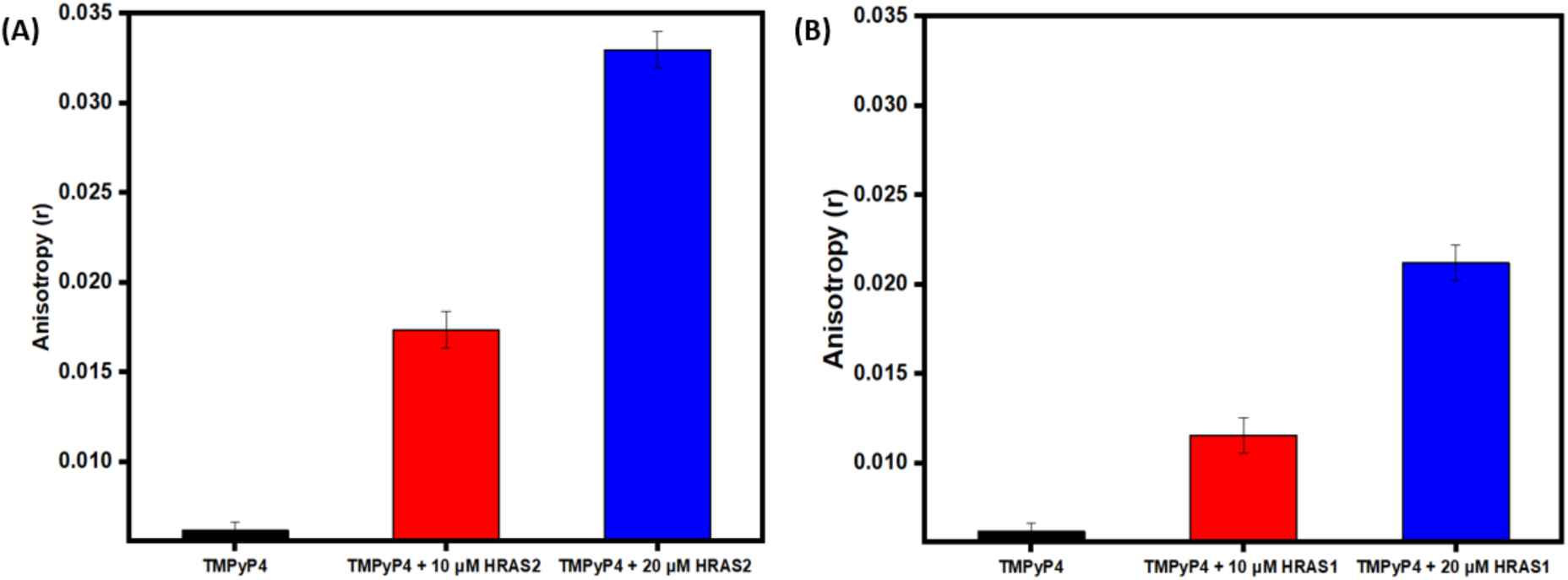
Fluorescence anisotropy decay (r) for the TMPyP4 (10 μM, λ_ex_ = 433 nm) in buffer and also in the HRAS2 (A) and HRAS1 (B) IM DNA (20 μM).

**Table 2:** Fluorescence anisotropy parameters of TMPyP4 (10 μM) in buffer and in the presence of different concentration (10 μM & 20 μM) of HRAS2 and HRAS1 iM DNA. *I_VV_, I_VH_, I_HV_* and *I_HH_* represent fluorescence signals for excitation and emission with the polarizer set at positions (0°,0°), (0°,90°), (90°,0°) and (90°,90°), respectively. G is the sensitivity factor of the detection systems (Temp-25^0^C; pH-5.4).

| *Sample* | *Anisotropy (r)* | *I<sub>HH</sub>* | *I<sub>HV</sub>* | *I<sub>VH</sub>* | *I<sub>VM</sub>* | *I<sub>VV</sub>* | *G Factor* |
| --- | --- | --- | --- | --- | --- | --- | --- |
| 10 μM TMPyP4 | 0.00617 | 163148.4 | 38313.4 | 85542.1 | 72809.1 | 20424.1 | 0.23 |
| 10 μM TMPyP4 + 10 μM HRAS2 iM | 0.01737 | 114799.6 | 27049 | 60175.33 | 51421.66 | 14801.66 | 0.23 |
| 10 μM TMPyP4 + 20 μM HRAS2 iM | 0.03299 | 81568.3 | 19340.7 | 42903.7 | 37176.7 | 11185.3 | 0.23 |
| 10 μM TMPyP4 + 10 μM HRAS1 iM | 0.01157 | 134571.9 | 31451.6 | 70869.96 | 60526.3 | 17167.6 | 0.23 |
| **10 μM TMPyP4**<br>**+ 20 μM HARS1**<br>**iM** | **0.02121** | 113327.<br>1 | 26606.<br>8 | 59630.4<br>6 | 51480.8 | 14968.<br>4 | 0.23 |

Typically, small molecules exhibit low anisotropy values in freely diffusing, low-viscosity environments. However, when such molecules associate with large, structured targets like DNA, proteins, membranes, or supramolecular assemblies (e.g., micelles or cyclodextrins), their rotational mobility is reduced, leading to elevated anisotropy signals. These distinctions make fluorescence anisotropy a powerful technique for examining binding modes and the dynamic constraints exerted by macromolecular complexes.

In buffer alone, TMPyP4 exhibited a baseline anisotropy value of 0.00617, which increased substantially to 0.03299 upon addition of 20 μM HRAS2 iM DNA (**Figure 4A**; **Table 1).** These pronounced increases in anisotropic values provide strong corroboration for earlier spectroscopic findings, supporting the conclusion that TMPyP4 bind to respective HRAS2 iM DNA. The elevated anisotropy values indicate reduced rotational freedom, suggesting that these compounds become embedded in more rigid, motionally restricted environments upon complex formation with the iM DNA.

### Time-Resolved Fluorescence Decay Reveals Microenvironmental Changes

To further validate and characterize the interactions of TMPyP4 with HRAS2 iM DNA structures, time-resolved fluorescence decay experiments were conducted [12]. Fluorescence lifetime analysis is a sensitive method for probing excited-state dynamics and provides insight into changes in the local microenvironment of fluorophores upon binding.

The average fluorescence lifetime (τ_avg_) was determined using tri-exponential decay fitting, allowing for a comprehensive evaluation of the overall emission behaviour rather than focusing on individual decay components [13]. Changes in τ_avg_ upon binding to HRAS2 iM DNA sequences were used to assess how the photophysical properties of TMPyP4 were influenced by these interactions. These lifetime shifts provide additional evidence for the formation of stable complexes and microenvironmental alterations upon binding to the HRAS2 iM DNA structures.

In the absence of HRAS2 iM DNA, the fluorescence decay of free TMPyP4 followed a tri-exponential model, characterized by a dominant component of 2.43 ns (95 % contribution), a long-lived component of 4.94 ns (4 %) and a very short-lived component of 0.20 ns (1 %), yielding an average lifetime (τ_avg_) of 2.52 ns. Upon binding to HRAS2 iM DNA, the decay profile also followed a tri-exponential pattern, consisting of a residual unbound fraction at 2.78 ns (5 %), a dominant component at 10.96 ns (78 %), and a long-lived component at 19.67 ns (17 %). The overall average lifetime increased to 11.13 ns **(Table 3**, **Figure 5A)**.

**Figure 5.**
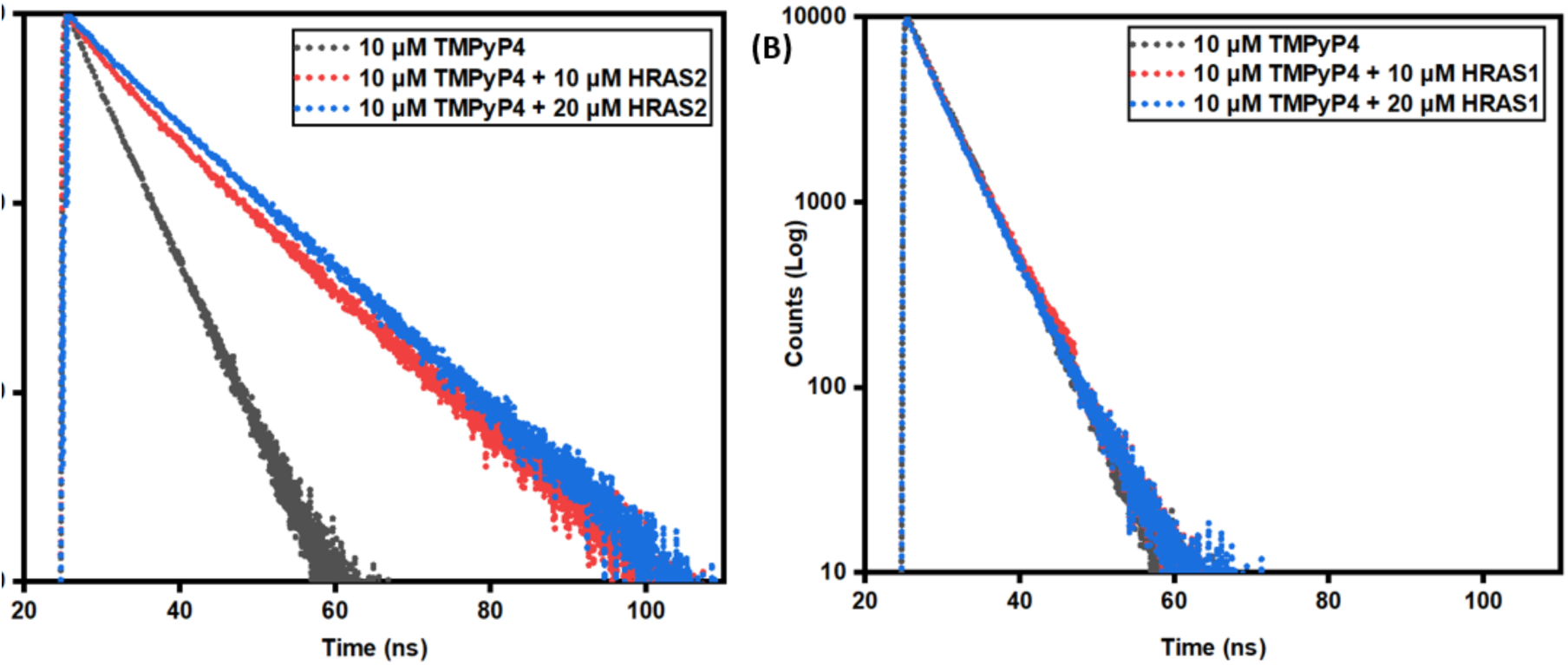
Time-Resolved fluorescence decay for the TMPyP4 (10 μM, λ_ex_ = 433 nm) in buffer and also in the HRAS2 (A) and HRAS1 (B) IM DNA (20 μM).

**Table 3:** Fluorescence decay parameters of TMPyP4 (10 μM) in buffer and in the presence of different concentration (10 μM & 20 μM) of HRAS2 and HRAS1 iM DNA.

| *Sample* | $\tau_1$<br>(ns) | $\alpha_1$ | $\tau_2$<br>(ns) | $\alpha_2$ | $\tau_3$<br>(ns) | $\alpha_3$ | $\tau_{avg}$<br>(ns) | $\chi^2$ |
| --- | --- | --- | --- | --- | --- | --- | --- | --- |
| **10 μM TMPyP4** | 2.43 | 0.95 | 4.94 | 0.04 | 0.20 | 0.01 | **2.52** | 1.02 |
| **10 μM TMPyP4**<br>**+ 10 μM HRAS2**<br>**iM** | 2.63 | 0.20 | 9.58 | 0.66 | 15.65 | 0.14 | **9.28** | 1.07 |
| **10 μM TMPyP4**<br>**+ 20 μM HRAS2**<br>**iM** | 2.78 | 0.05 | 10.96 | 0.78 | 19.67 | 0.17 | **11.13** | 1.06 |
| **10 μM TMPyP4**<br>**+ 10 μM HRAS1**<br>**iM** | 0.37 | 0.03 | 4.42 | 0.66 | 6.39 | 0.31 | **3.72** | 1.04 |
| **10 μM TMPyP4**<br>**+ 20 μM HARS1**<br>**iM** | 0.36 | 0.03 | 4.45 | 0.67 | 6.41 | 0.30 | **3.74** | 1.03 |

The bound forms of TMPyP4 exhibited longer fluorescence lifetimes than their unbound states. This extension of lifetime suggests a transition into less polar microenvironments upon iM DNA binding. Specifically, the interaction of TMPyP4 with HRAS2 likely positions the molecule within hydrophobic regions of the iM DNA structure. The restricted access to water in these domains reduces non-radiative decay pathways, stabilizing the excited states and increasing fluorescence lifetime.

This behaviour is consistent with the hydrophobic nature of the core scaffolds of TMPyP4. Binding likely involves de-solvation from the aqueous buffer into the more hydrophobic iM DNA interior, providing a thermodynamic driving force for complex formation. Interestingly, the binding selectivity of TMPyP4 appears to be influenced by its structural features. Based on our experimental findings, it may be predicted that the flat porphyrin core along with pyridyl groups contributes to increased polarity, especially upon protonation, which could favour interaction with more exposed or polar iM regions, such as those present in HRAS2.

### Circular Dichroism Spectroscopy Indicates Structural Alterations of HRAS2 iM DNA Upon TMPyP4 Binding

Circular dichroism (CD) spectroscopy is a sensitive technique widely used to monitor conformational features and backbone dynamics of iM structures, and it has proven particularly effective for characterizing DNA–ligand interactions [12]. In this study, CD analysis was employed to probe the secondary structures of HRAS2 and HRAS1 iM DNA sequences, and to assess any conformational changes resulting from their interaction with TMPyP4.

In the absence of TMPyP4, the CD spectra of HRAS2 and HRAS1 iM DNAs displayed distinct and characteristic signatures: HRAS2 presented a positive peak at 290 nm and a negative peak at 258 nm **(Figure 6A)**; while HRAS1 showed a positive band at 278 nm and a negative band at 242 nm **(Figure 6B)**. These spectral features are consistent with established profiles of well-folded iM DNA structures.

**Figure 6.**
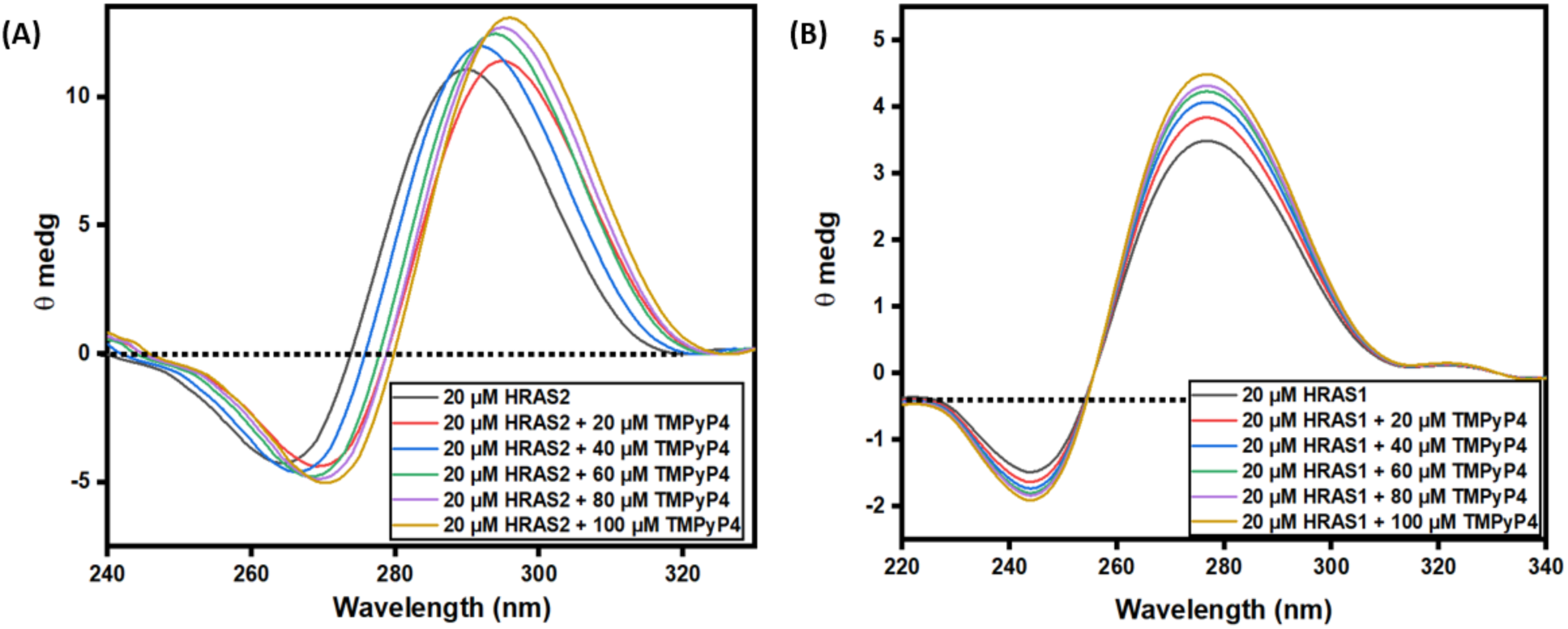
Circular dichroic spectral profile of HRAS2 (A) and HRAS1 (B) IM DNA (20μM) in the presence of TMPyP4 (20 μM, 40 μM, 60 μM, 80 μM, 100 μM).

Upon titration with increasing concentrations of TMPyP4, significant alterations in the intensity and wavelength positions of both the positive and negative CD bands of HRAS2 were observed. These significant spectral variations suggest that the core HRAS2 iM topology is substantially perturbed upon TMPyP4 binding. The presence of significant shifts or alterations in the HRAS2 CD signatures implies that TMPyP4 interacts with HRAS2 iM DNA, leading to disruption of its folded conformation. In contrast, no significant changes in the CD profile of HRAS1 were observed upon addition of TMPyP4.

### UV Melting Studies Reveal Enhanced Thermal Stability of HRAS2 iM DNA Structures Upon TMPyP4 Binding

Thermal denaturation, or UV melting analysis, offers a robust method for evaluating the stabilizing effects of small molecules on iM DNA structures [12]. It is well established that ligands engaging in base-stacking interactions with iM DNA can increase the melting temperature (T_m_) by enhancing the structural stability of the folded structure. In contrast, ligands that bind via electrostatic or groove interactions often induce minimal or no change in T_m_ . To assess the impact of TMPyP4 on the thermal stability of HRAS2 and HRAS1 iM DNA, we conducted UV melting experiments by monitoring absorbance changes at 295 nm, a wavelength characteristic of iM melting transitions. Tm values were determined from the the midpoint (inflection point) of each melting curve.

As shown in **Figure 7A**, the addition of 20 μM TMPyP4 led to a clear increase in the T_m_ of the HRAS2 iM DNA target. This significant thermal shift indicates enhanced stabilization of the iM fold upon ligand binding.

**Figure 7.**
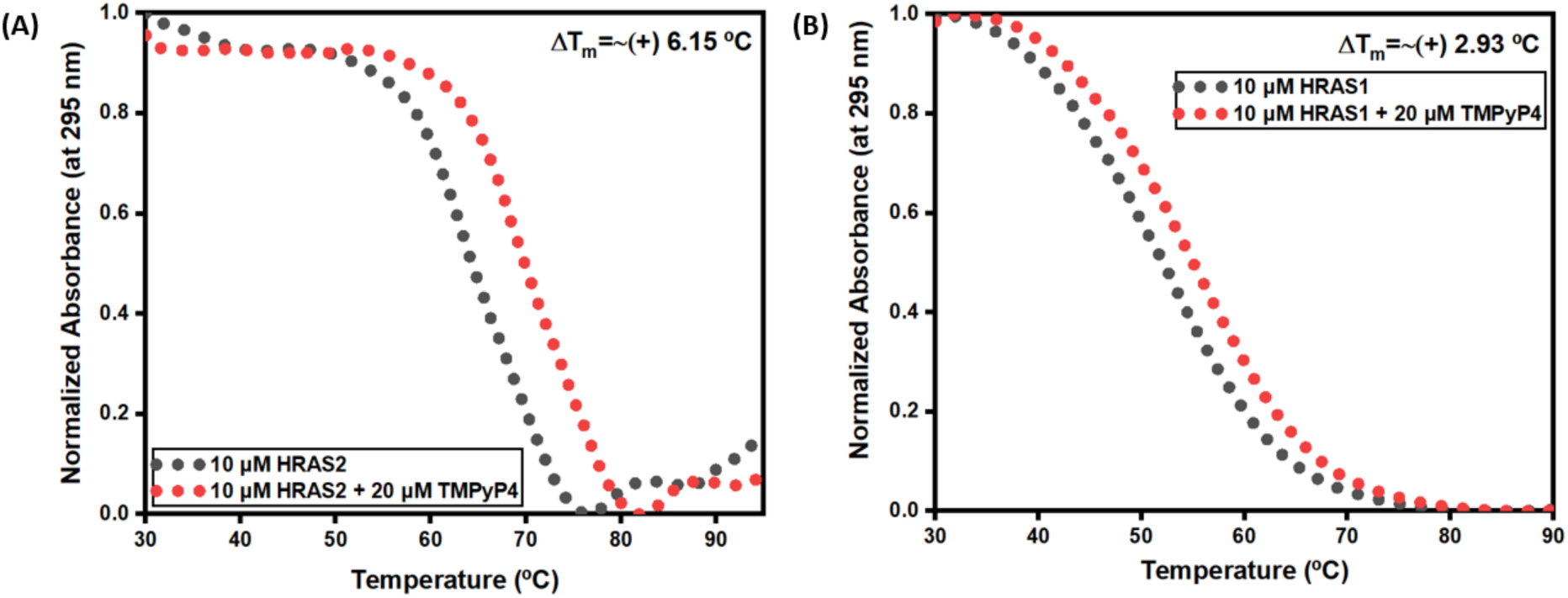
Thermal melting profiles of HRAS2 (A) and HRAS1 (B) IM DNA (10 μM) alone and in the presence of TMPyP4 (20 μM).

The observed T_m_ increases support a binding mechanism dominated by base-stacking interactions between TMPyP4and the HRAS2 iM DNA core. These findings are consistent with our spectroscopic and fluorescence lifetime data, collectively supporting that TMPyP4 engages in a stabilizing, high-affinity interaction with the HRAS2 iM DNA structure. Notably, the extent of thermal stabilization differs between HRAS2 and HRAS1 iM sequences, suggesting a degree of structural selectivity in TMPyP4 binding.

From the UV melting studies, the thermodynamic parameters (ΔH, TΔS, and ΔG) were determined for the interaction between TMPyP4 and HRAS2 iM DNA using Van’t Hoff analysis, and the associated errors were calculated accordingly. Curve fitting was performed through nonlinear regression to ensure the accuracy and reliability of the obtained values [15, 16]. At a concentration of 20 μM TMPyP4, the enthalpy change (ΔH = −64.91 kcal mol⁻¹) was notably larger in magnitude than the entropy contribution (TΔS = −55.69 kcal mol⁻¹). This suggests that the binding process is predominantly governed by enthalpic factors. The negative Gibbs free energy (ΔG = −9.22 kcal mol⁻¹) further indicates that the interaction occurs spontaneously under the studied conditions **(Table 4).**

**Table 4:** Thermodynamic parameters for 10 μM HRAS2 iM DNA in the absence and in the presence of 20 μM TMPyP4. Thermal melting curves were normalized and analysed through curve fitting using Kaleida Graph (Synergy Software).

| *Sample* | *Temp*<br>(°C) | *ΔH*<br>(kcal/mol) | *TΔS(kcal/mol)* | *ΔG*<br>(kcal/mol) |
| --- | --- | --- | --- | --- |
| **10 μM HRAS2 iM** | 25 | -53.42±0.2 | -50.89±0.2 | -2.53±0.1 |
| **10 μM HRAS2 iM + 20 μM**<br>**TMPyP4** | 25 | -64.91±0.1 | -55.69±0.1 | -9.22±0.2 |

Overall, the consistently large negative enthalpy values, compared to the relatively smaller entropy contributions, indicate that the binding process is mainly enthalpy-driven. This points toward the involvement of strong and specific interactions between the ligand TMPyP4 and the HRAS2 iM DNA. Although entropy does contribute to the process, its role appears secondary, suggesting that the stability of these complexes is largely controlled by favourable enthalpic interactions.

### FT-IR Spectroscopy Reveals Ligand-Induced Structural Changes in iM DNA

Fourier Transform Infrared (FT-IR) spectroscopy is a powerful analytical technique that utilizes infrared radiation to probe molecular vibrations and assess structural features of biomolecules. In the context of cytosine-rich DNA, FT-IR has been widely applied to investigate structural characteristics and conformational changes associated with iM DNA formation, providing insights into ligand-induced perturbations [17,18].

In this study, FT-IR spectroscopy was employed to examine potential structural perturbations in HRAS2 and HRAS1 iM DNA sequences upon binding to TMPyP4. The spectral region from 900 to 2000 cm⁻¹ was analysed, as it contains characteristic vibrational bands attributed to in-plane C=C ring stretching and carbonyl group vibrations of the nucleobases.

As illustrated in **Figure 8**, the FT-IR spectra exhibit distinct peaks in the 800–2000 cm⁻¹ range, indicating that the iM DNA structures are retained under the hydration conditions used. This spectral window includes characteristic vibrational bands attributed to multiple structural components of DNA, such as the sugar backbone, phosphate groups, and nucleobases. FT-IR spectroscopy is a versatile technique capable of probing DNA conformation in various states, including liquids, solids, gels, and fibres. In D₂O, the 1500–1800 cm⁻¹ region is particularly responsive to base-specific vibrational modes, including in-plane bending and stretching of double bonds such as C=N, C=C, and C=O. These bands are sensitive to alterations in base stacking, hydrogen bonding, hydration dynamics, and metal ion coordination, due to the electron-rich nature of π-bond systems. Variations in spectral position and band intensity across this region reflect both conformational and torsional changes within the DNA structure.

**Figure 8.**
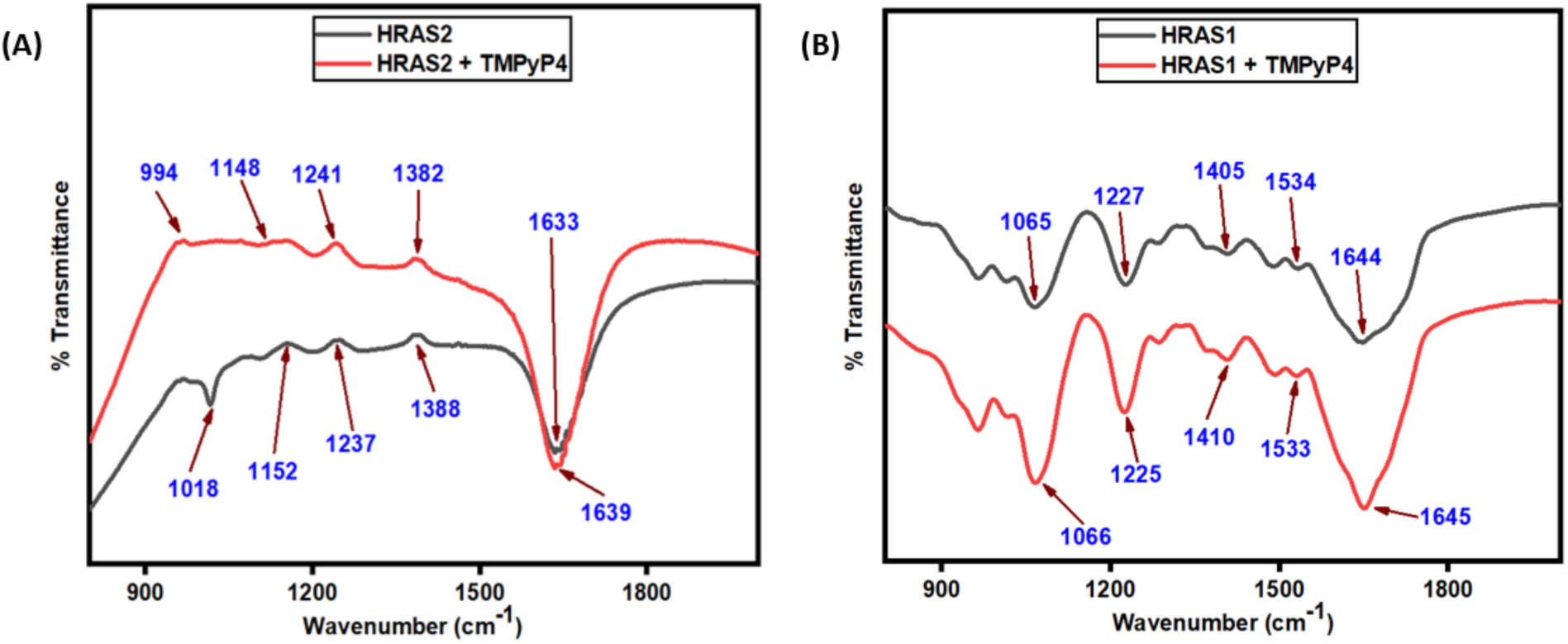
Fourier Transform Infrared (FT-IR) spectra in the [800–2000] cm^-1^ spectral region of HRAS2 IM DNA (A, black line), HRAS1 IM DNA (B, black line) and HRAS2 IM DNA+ TMPyP4 (A, red line), HRAS1 IM DNA+ TMPyP4 (B, red line). The peak centre of each spectrum is indicated by the arrows.

**Figures 8A–8B** display the FT-IR spectra of HRAS2 and HRAS1 iM DNAs, confirming the retention of their folded structures under the tested hydration conditions. Upon addition of TMPyP4, notable spectral shifts and alterations in peak positions were observed in HRAS2 iM DNA. These shifts reflect changes in the local environment of the nucleobases, suggesting that ligand binding may induce conformational rearrangements within the HARS2 iM structure. Overall, the FT-IR data support the presence of ligand-induced structural perturbations in HRAS2 iM DNA, highlighting the dynamic nature of these complexes and further corroborating the interaction between TMPyP4 and HRAS2 iM DNA.

## CONCLUSIONS

In this work, we examined how TMPyP4 interacts with several iM-forming DNA sequences that originate from the promoter regions of cancer-related genes, including HRAS1, HRAS2, BCL2, CMYC, CKIT, VEGF, and the human telomeric sequence. A range of spectroscopic and biophysical techniques was used to understand these interactions in detail. By combining UV–Vis absorption, steady-state and time-resolved fluorescence, fluorescence anisotropy, circular dichroism (CD), thermal melting studies, and FT-IR spectroscopy, we were able to evaluate both the binding behaviour of TMPyP4 and the structural changes that occur in the DNA upon ligand association.

Among the different i-motif sequences examined, HRAS2 iM DNA showed the strongest response in the presence of TMPyP4, indicating a higher binding preference compared with the other tested sequences. The spectroscopic experiments consistently pointed toward the formation of a stable interaction between TMPyP4 and the HRAS2 iM structure. Changes observed in absorption and fluorescence measurements suggested that TMPyP4 associates closely with the HRAS2 iM DNA structure, while fluorescence lifetime and anisotropy results further supported the formation of a stable TMPyP4–HRAS2 iM DNA complex.

Structural analysis also provided important information about the effect of TMPyP4 on the HRAS2 iM conformation. Circular dichroism measurements indicated that the overall HRAS2 iM topology was altered upon binding, suggesting that TMPyP4 stabilizes the structure rather than disrupting it. This stabilizing effect was further confirmed by thermal melting experiments, where the HRAS2 iM displayed improved thermal stability in the presence of TMPyP4. Based on the thermodynamic results, the binding of TMPyP4 to HRAS2 iM DNA appears to be mainly enthalpy-driven, with the process proceeding spontaneously under the studied conditions. Additional insights from FT-IR spectroscopy indicated subtle local changes in the HRAS2 iM DNA environment after binding, pointing to specific molecular interactions between TMPyP4 and the cytosine-rich HRAS2 iM DNA sequence.

The interaction between TMPyP4 and HRAS2 iM DNA can be explained by the complementary features of the molecule and the DNA structure. TMPyP4 has a flat aromatic core and carries positive charges, which makes it naturally attracted to DNA, whose backbone is negatively charged. This attraction helps the molecule approach and associate with the DNA strand. In addition, the flat structure of TMPyP4 allows it to align with the stacked bases present in the HRAS2 iM, helping to stabilize the structure. Variations in the sequence and folding pattern of different iMs can also influence the strength of binding, which may explain why certain sequences, such as HRAS2, show stronger interactions than others.

Taken together, these results show that TMPyP4 can selectively recognize and stabilize the HRAS2 iM structure among several cancer-associated iM sequences. While TMPyP4 has been widely investigated for its interaction with G-quadruplex DNA, studies focusing on its behaviour toward iM structures are still relatively limited. The findings of this work therefore provide useful information about how small molecules interact with cytosine-rich DNA motifs. In addition, the spectroscopic response observed upon binding suggests that TMPyP4 may also serve as a practical fluorescent probe for studying iM structures. Overall, this study contributes to the growing interest in non-canonical DNA structures as potential molecular targets in cancer-related research. A better understanding of how these molecules interact with such DNA motifs may help in the future development of selective probes and therapeutic agents designed to regulate gene expression through iM targeting.

## Notes

The authors declare no competing financial interest.

## ACKNOWLEDGMENTS

Sudipta Bhowmik thanks “Intramural Seed Money Research Committee, SBV” for “SBV-Seed money” (SBV/IRC/SEED MONEY/167/2023) for necessary research chemicals & consumables.

## References

1. Chu, C., Zhang, S., Guan, Z., Shen, J., Luo, T., Zhou, J., Mergny, J.L. and Cheng, M., 2026. High-throughput measurement and prediction of the i-motif DNA stability landscape. Nucleic Acids Research, 54(4), p.gkag110.

2. Adala, J.D. and Knutson, B.A., 2026. In silico mapping of non-canonical DNA structures across the human ribosomal DNA locus. G3: Genes, Genomes, Genetics, 16(2), p.jkaf299.

3. Bochalis, E., Dereki, I., Wang, G., Sgourou, A., Vasquez, K.M. and Georgakopoulos-Soares, I., 2026. Non-B DNA structures and their contributions to genetic diversity, aging, and disease. Nucleic Acids Research, 54(4), p.gkag084.

4. Sengupta, P., Gillet, N., Obi, I. and Sabouri, N., 2026. Mechanistic insights into PCBP1-driven unfolding of selected i-motif DNA at G1/S checkpoint. Nature Communications, 17(1), p.1149.

5. Hamer, M., Argañaras, O.J. and Narambuena, C.F., 2026. Minimal requirements for one-dimensional aggregation in simple coarse-grained models of charged porphyrinoid units. Physical Chemistry Chemical Physics, 28(9), pp.5796–5806.

6. Sokulska, M., Nalewaj, M., Czapik, T., Rachwalak, M. and Szabat, M., 2026. Modulating G-quadruplexes for therapeutic intervention: structural diversity, stability, and emerging nucleic acid–based strategies. Molecular Therapy Nucleic Acids.

7. Li, P., Zhou, D., Xie, Y., Yuan, Z., Huang, M., Xu, G., Huang, J., Zhuang, Z., Luo, Y., Yu, H. and Wang, X., 2024. Targeting G-quadruplex by TMPyP4 for inhibition of colorectal cancer through cell cycle arrest and boosting anti-tumor immunity. Cell Death & Disease, 15(11), p.816.

8. Iachettini, S., Biroccio, A. and Zizza, P., 2024. Therapeutic use of G4-ligands in cancer: state-of-the-art and future perspectives. Pharmaceuticals, 17(6), p.771.

9. Alici, E.H., Bilgicli, A.T., Tüzün, B., Günsel, A., Arabaci, G. and Yarasir, M.N., 2022. Alkyl chain modified metalophthalocyanines with enhanced antioxidant-antimicrobial properties by doping Ag+ and Pd2+ ions. Journal of Molecular Structure, 1257, p.132634.

10. Bag, S., Chand, K., Burman, M.D., Vertueux, S., Chorell, E. and Bhowmik, S., 2025. Exploring i-Motif DNA binding with benzothiazolino Coumarins: Synthesis, Screening, and spectroscopic insights. Bioorganic chemistry, 156, p.108227.

11. Bag, S., Ghosal, S., Mukherjee, M., Pramanik, G. and Bhowmik, S., 2024. Quercetin exhibits preferential binding interaction by selectively targeting HRAS1 I-Motif DNA-forming promoter sequences. Langmuir, 40(19), pp.10157–10170.

12. Bag, S., Ghosal, S., Karmakar, S., Pramanik, G. and Bhowmik, S., 2023. Uncovering the contrasting binding behavior of plant flavonoids fisetin and morin having subsidiary hydroxyl groups (− OH) with HRAS1 and HRAS2 i-Motif DNA structures: decoding the structural alterations and positional influences. ACS omega, 8(33), pp.30315–30329.

13. Lisitsyna, E.S., Klose, A., Vuorimaa-Laukkanen, E., Ijäs, H., Lajunen, T., Suhling, K., Linko, V. and Laaksonen, T., 2025. Fluorescence anisotropy for detailed analysis of doxorubicin loading into dna origami nanocarriers for drug delivery. ACS Applied Nano Materials, 8(26), pp.13274–13284.

14. Bag, S. and Bhowmik, S., 2023. Fluorescence spectroscopy: A useful method to explore the interactions of small molecule ligands with DNA structures. In Reverse Engineering of Regulatory Networks (pp. 33–49). New York, NY: Springer US.

15. Takahashi, S., Bhattacharjee, S., Ghosh, S., Sugimoto, N. and Bhowmik, S., 2020. Preferential targeting cancer-related i-motif DNAs by the plant flavonol fisetin for theranostics applications. Scientific Reports, 10(1), p.2504.

16. Takahashi, S. and Sugimoto, N., 2017. Quantitative Analysis of Nucleic Acid Stability with Ligands Under High Pressure to Design Novel Drugs Targeting G-Quadruplexes. Current Protocols in Nucleic Acid Chemistry, 70(1), pp.17–9.

17. Bag, S., Ghosal, S., Burman, M.D., Manna, A. and Bhowmik, S., 2025. Structural Diversity and Mechanistic Insights on Preferential Interaction of Small Molecule Ligands with i-Motif DNA Structures: Unlocking New Blueprint for Drug Discovery. ChemistrySelect, 10(24), p.e01985.

18. Bhusari, N., Bagul, A., Mishra, V.K., Tufail, A., Gaikwad, D. and Dubey, A., 2025. Sustainable synthesis of Schiff base derivatives via an ionic liquid and a microwave-assisted approach: structural, biological, and computational evaluation. RSC advances, 15(28), pp.22764–22788.

