## Supporting Information for "Selective Stabilization of HRAS2 i-Motif DNA by TMPyP4: A Multimodal Biophysical and Thermodynamic Investigation"

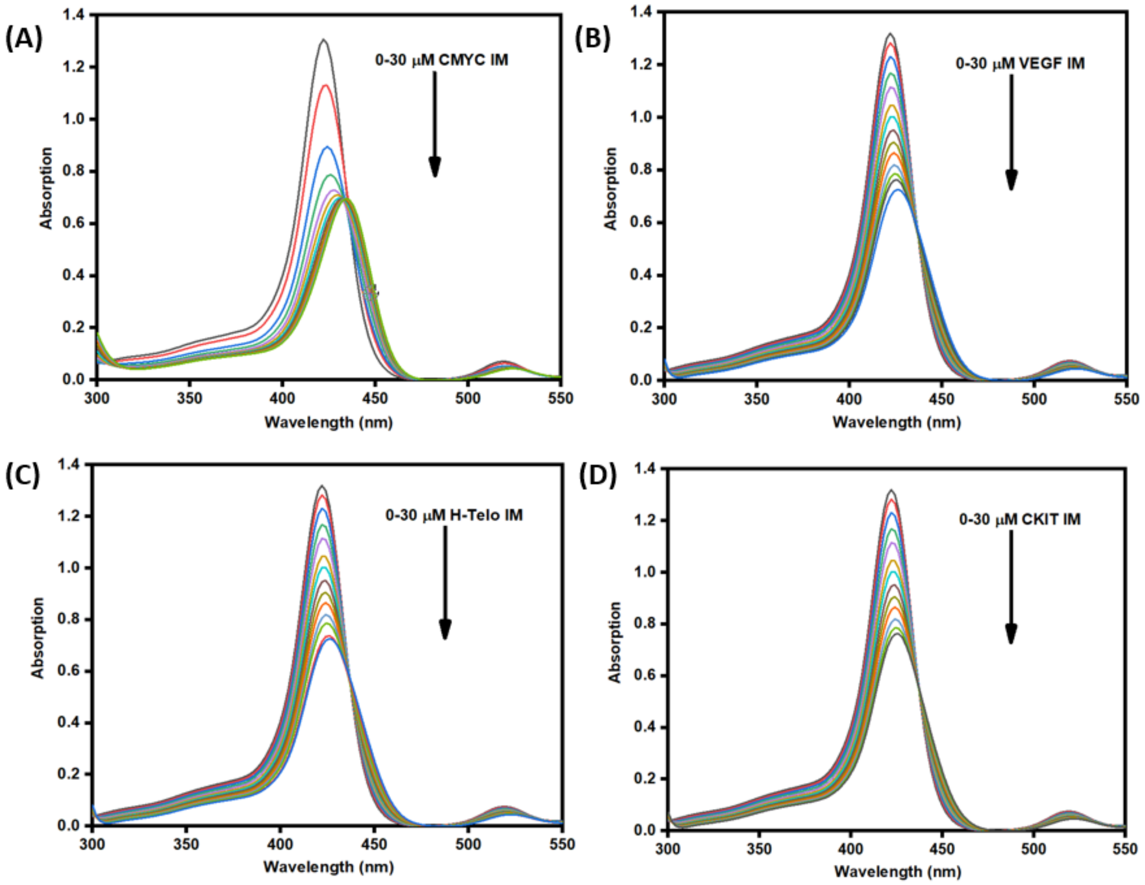


**Figure S1.** UV-Absorption emission spectra of TMPyP4 (15 μM) in the absence and presence of successive additions of CMYC (A), VEGF (B), H-Telo (C) and CKIT (D) IM DNA (30 μM).

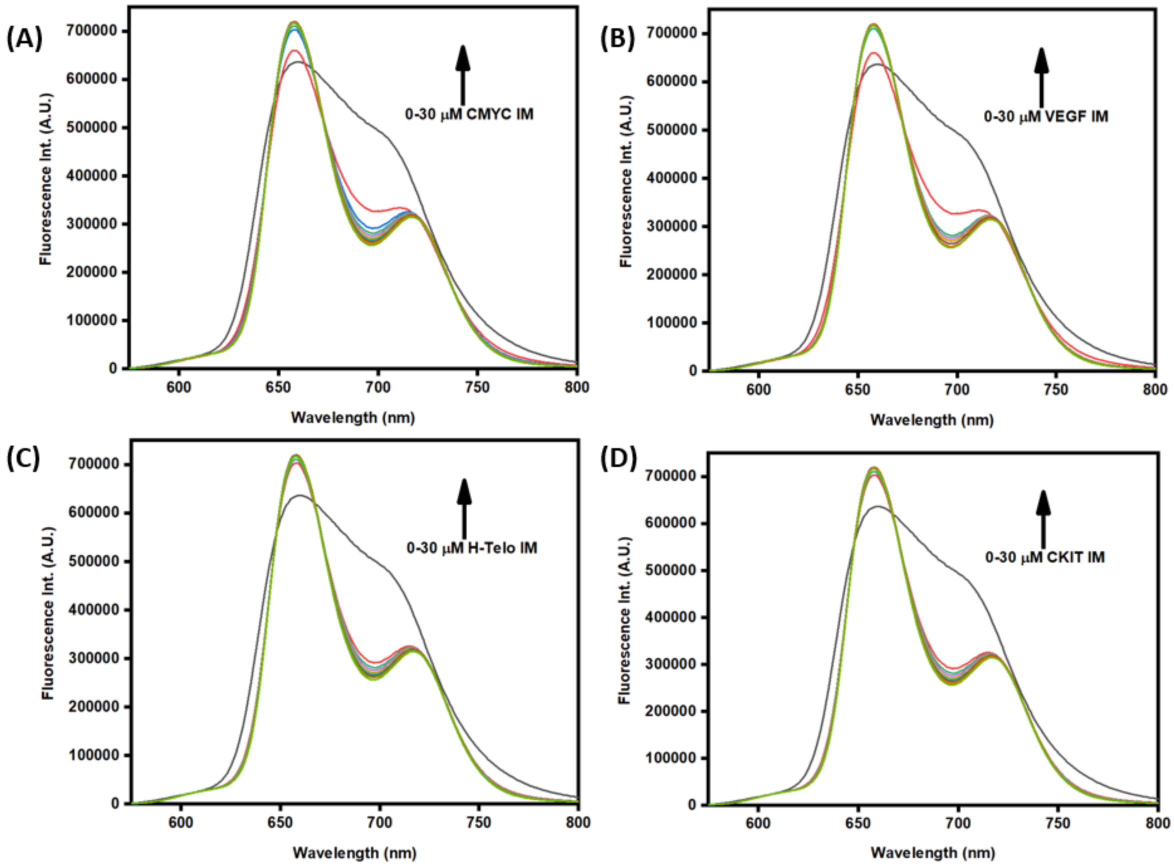


**Figure S2.** Steady state fluorescence emission spectra of TMPyP4 (15 μM) in the absence and presence of successive additions of CMYC (A), VEGF (B), H-Telo (C) and CKIT (D) IM DNA (30 μM).
